# The Aggregated Gut Viral Catalogue (AVrC): A Unified Resource for Exploring the Viral Diversity of the Human Gut

**DOI:** 10.1101/2024.06.24.600367

**Authors:** Anastasia Galperina, Gabriele Andrea Lugli, Christian Milani, Willem M. De Vos, Marco Ventura, Anne Salonen, Bonnie Hurwitz, Alise J. Ponsero

## Abstract

Despite the growing interest in the role of the gut virome in human health and disease, identifying viral sequences from human gut metagenomes remains computationally challenging due to underrepresentation of viral genomes in reference databases. Several recent large-scale efforts have mined human gut metagenomes to establish viral sequence catalogues, using varied computational tools and quality control criteria. However, there has been no consistent comparison of these catalogues’ quality, diversity, and completeness, nor unification into a comprehensive resource. Here, we systematically surveyed nine previously published human gut viral catalogues, assessing their quality and the overlap of the viral sequences retrieved. While these catalogues collectively screened >40,000 human fecal metagenomes, 82% of the recovered 345,613 viral sequences were unique to one catalogue, highlighting limited redundancy. We further expanded representation by mining 7,867 infant gut metagenomes, retrieving 1,205,739 additional putative viral sequences. From these datasets, we constructed the Aggregated Gut Viral Catalogue (AVrC), a unified modular resource containing 1,018,941 dereplicated viral sequences (449,859 species-level vOTUs). Detailed annotations were generated for sequence quality, taxonomy, predicted lifestyle, and putative host. The AVrC reveals the gut virome’s substantial unexplored diversity, providing a pivotal resource for viral discovery. The AVrC is accessible as a relational database and through a web interface allowing customized querying and subset retrieval, enabling streamlined utilization by the research community and future expansions as novel data becomes available.

**Author summary:** The human gut is home to a vast array of viruses, collectively known as the gut virome, which play a crucial role in human health and disease. Recently, several research groups aiming at providing an overview of the Human gut viral diversity, have created catalogues of viral sequences found in the human gut by analyzing a large number of fecal samples from different individuals. In this study, we compared nine of these existing catalogues and found that there was surprisingly little overlap between them, with 82% of the viral sequences being unique to a single catalogue. To further expand the available data, we analyzed nearly 8,000 additional fecal samples from infants. By combining all this ressources, we created a unified resource called the Aggregated Gut Viral Catalogue (AVrC), which contains more than a million distinct viral sequences, representing nearly 450,000 different viral species. This catalogue, which is easily accessible to the scientific community through a user-friendly web interface, provides a valuable tool for exploring the vast diversity of the human gut virome and its potential implications for human health.

## Introduction

Despite the increasing number of studies highlighting the importance of gut virome in health and disease, identifying viral sequences in large metagenomic datasets is still computationally challenging. Strikingly, in gut viromes, 75% to 99% of viral reads do not produce significant alignments to any known viral genome (1). This large range can be partially explained by broad under-representation of viral sequences in most genomic databases and the overrepresentation of specific virus taxonomic groups in these databases. All in all, gut virome profiling based on viral RefSeq databases is shown to lead to a poor and incomplete delineation of the true gut virome composition (2).

Recent bioinformatic tools leverage machine learning algorithms to identify features that signal a phage origin, and typically allow for a broader recall of previously unknown sequences than reference-based approaches. VirFinder (3), DeepVirFinder (4) and Seeker (5) use a machine-learning approach to classify sequences as phage or prokaryotic based on their k-mer sequence composition. Other machine-learning tools such as VirSorter2 (6), VirMiner (7), VIBRANT (8), base their prediction on genomic features such as the relative synonymous codon usage, gene density, strand shifts, and the number of protein gene homologs. Altogether, these new approaches provide new avenues to explore the untapped viral diversity in metagenomes. However, all currently available tools are limited to the classification of assembled contigs (cannot be applied to raw sequencing reads), and typically provide a simple binary classification (viral vs non-viral).

In the past few years, several large-scale aggregation efforts mining viral sequences from metagenomes have been released. The IMG/vr database maintained by the Department of Energy (DOE) (9,10) mines both environmental and human-associated metagenomes, while several viral catalogues focus specifically on the human gut: Gregory et al. analyzed 2,697 human gut metagenomes, recovered 33,242 species-like viral operational taxonomic units (vOTUs), and established the gut virome database (GVD) (11). Tisza et al. collected 5,996 metagenome datasets from human gut, skin, saliva and vagina to build the Cenote human virome database (CHVD) containing 45,033 species-like vOTUs (12). Benler et al. mined the human gut metagenomes available in NCBI in 2019 and retrieved 3738 complete phage genomes (13), Nishijima et al. explored the viral diversity associated to the gut microbiota of healthy and diseased adult Japanese (14) and Van Espen et al. surveyed the gut virome associated with 91 healthy Danish children, adolescents, and adults (15). Nayfach et al. constructed the metagenomic gut virus (MGV) catalog from 11,810 human gut metagenomes and retrieved 54,118 species-like vOTUs (16). To date, the largest effort is the mining of 28,060 human gut metagenomes by Camarillo-Guerrero et al. to generate the gut phage database (GPD), which contained 142,809 species-like vOTUs (17). Recently, some efforts focused on infant gut metagenomes, for example the COPSAC virome dataset leveraged a dataset of 465 infant fecal metagenomes and retrieved 10,021 vOTUs (18).

Despite the value of these large-scale efforts in the human gut virome, there is currently no consistent comparison of the quality, diversity, and completeness of these catalogues. Additionally, there is a currently unmet need to aggregate these viral sequences in a cohesive resource, that users can leverage to compare their own viral sequences and that can be expended as novel viral sequences are published. Here, we aimed to (1) survey previous viral mining efforts in human fecal metagenomes, assess their quality, diversity and overlap; and to (2) harmonize and aggregate all currently available gut viral catalogues in a unified resource that users can leverage to compare new putative viral sequences to previous mining efforts.

## Results

### 1. Previous efforts to generate human gut viral catalogues have disparate quality and a limited overlap

We identified eight studies all aiming to generate a catalogue of viral sequences derived from human gut metagenomes, published between 2020 and 2023 (**Table 1**). Additionally, the IMG/Vr database is a dedicated resource developed specifically to retrieve viral sequences from metagenomes made available through the DOE JGI service and contains a number of human gut derived viral sequences. The tools and pipelines used for the identification and retrieval of viral sequence were highly different across studies. Almost all used one or several dedicated viral identification tools, that perform a classification based on k-mer composition (VirFinder, DeepVirFinder, Seeker) or on the identification of genomic features (VirSorter, ViralVerify, geNomad). Only the COPSAC infant study used a completely custom viral identification method. Most of studies confirmed the viral origin and evaluated the quality of the retrieved sequences using the dedicated tool CheckV (**Table 1**). Although sample-associated metadata could not be retrieved for all the published studies, we estimate that altogether, more than 40,000 human fecal samples from 40 different countries were screened to generate these eight catalogues. Importantly, we estimate that 40% of the metagenomes previously screened for viral sequences were generated from healthy adult stool samples, and that at least 30% of the samples were mined in more than one catalogue (**Supplemental File 1**). Most catalogues screened bulk fecal metagenomes, and only the COPSAC and the DEVoC exclusively included viromes from VLP-enriched fecal samples. The GVD screened metagenomes of both VLP-enriched and bulk metagenomes.

**Table 1:** List of public viral catalogues from human gut metagenomes and details of the size and computational method used to generate the catalogues. * Subset of vOTUs from “Human gut” ecosystem accessed July 2023 ** Subset of vOTUs signaled as present in gut metagenomes

| Source name | Source type | Reference | Viral Identification tool | QC |
| --- | --- | --- | --- | --- |
| *IMG/Vr* | Metagenomic database | (10) | geNomad | CheckV |
| *Gut virome database (GVD)* | Metagenomic catalogue of human gut virome | (11) | VirSorter<br>VirFinder<br>CAT | BUSCO<br>hmmsearch |
| *Gut Phage database (GPD)* | Metagenomic catalogue of human gut virome | (17) | VirSorter<br>VirFinder | Custom ML model<br>CheckV |
| *Metagenomic gut virus (MGV)* | Metagenomic catalogue of human gut virome | (16) | Custom protein search<br>VirFinder<br>VirSorter | CheckV |
| *Cenote human virome database (CHVD)* | Metagenomes from adult gut, skin, vaginal and oral. | (12) | Cenote-Taker 2 | CheckV |
| *COPSAC infant phages* | Metagenomes from infant from the COPSAC cohort | (18) | Protein based custom identification | Custom QC |
| *Gut Phages (KGP)* | Metagenomes from adult gut virome | (13) | Protein based identification<br>Seeker<br>ViralVerify | Custom QC |
| *Japanese 4D catalogue* | Metagenomes from adult gut virome | (14) | DeepVirFinder<br>HMM-HMM<br>fetchMG<br>barrnap<br>DIAMOND | CheckV |
| *Danish Enteric Virome Catalog (DEVoC)* | Metagenomes from children, adolescents, and adults | (15) | VirSorter<br>Database search<br>Gene content and structure | CheckV |

While all pipelines used to generate each catalogue contained a quality control step, the overall quality of the sequences as evaluated by CheckV where highly variable with catalogues such as DEVoC composed of more than 75% of low-quality sequences, while others, such as the MGV, KGP, or Japanese 4D, containing less than 5% of low-quality sequences (**Fig. 1A**). As some viral identification tools can be biased toward a misclassification of plasmids as viral sequences, we assessed the potential plasmid contamination of each catalogue using geNomad. Most catalogues contained some plasmid contamination, with the GVD containing more than 10% of sequences potentially arising from plasmids. Notably, the recent release of the IMG/Vr database includes a plasmid decontamination step, and therefore did not contain any detectable plasmid contamination (**Fig. 1A**).

**Fig 1:**
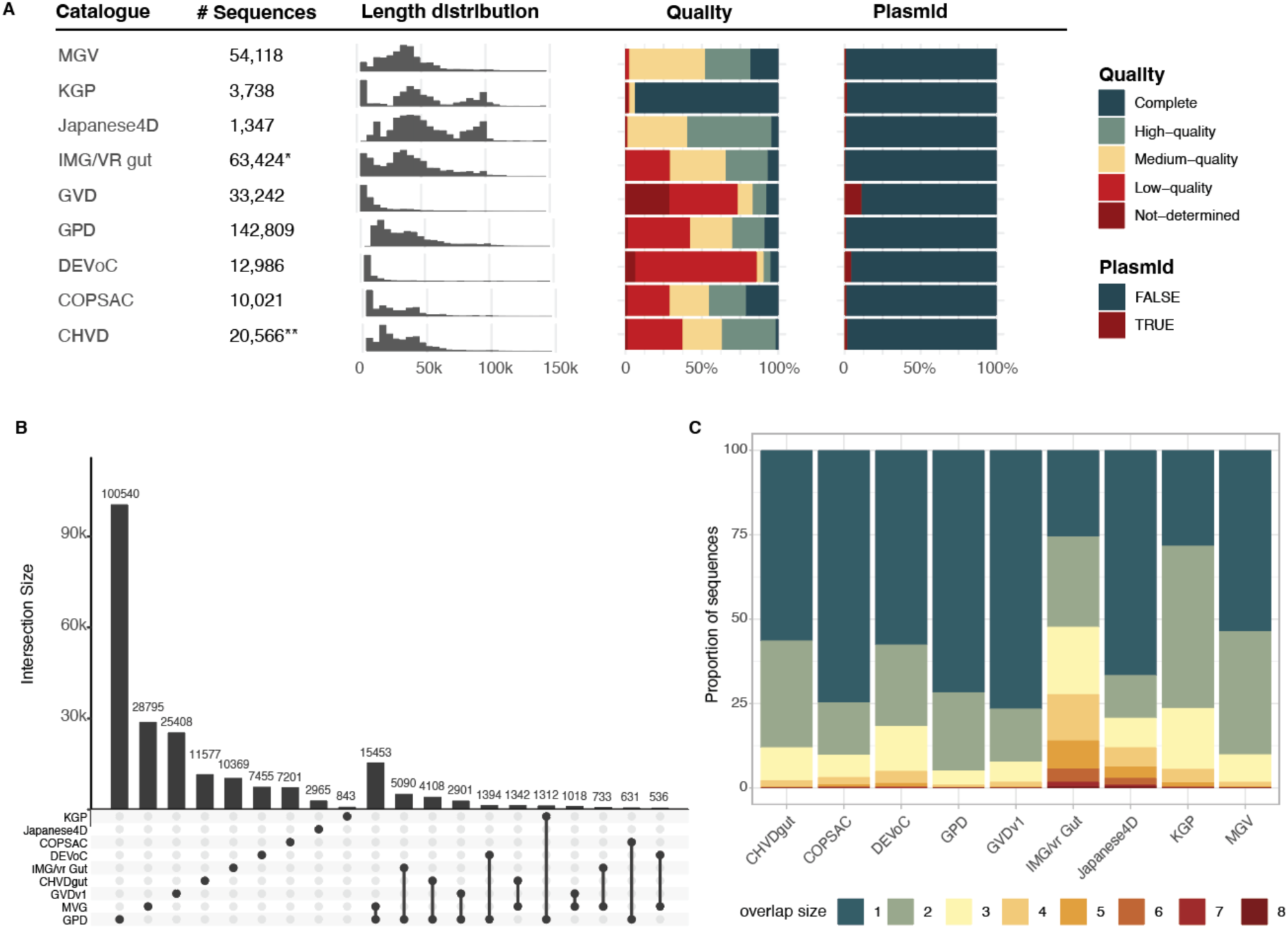
Overview of previously published gut viral catalogues. A: Previously published catalogues size, viral sequence length distribution, viral sequences quality and potential plasmid contamination. B: UpSet plot of the vOTU overlap between the previously published catalogues. The viral sequences from all the catalogues were clustered into vOTU and shared vOTU are defined as a cluster that grouped sequences from different catalogues. The intersection size was computed as the number of vOTU shared by the catalogues. The columns are sorted based on the vOTU counts per catalogue and their overlap between all combinations of catalogues. C: Proportion of unique and shared vOTU in the previously published catalogues. The sequences in the catalogues were clustered into vOTU and the “overlap size” of each vOTU was defined as the number of catalogues that contained at least one sequence for that vOTU. An overlap size of one signifies that the vOTU was uniquely found in the considered catalogue. * Subset of vOTUs from “Human gut” ecosystem accessed July 2023 ** Subset of vOTUs signaled as present in gut metagenomes

We next evaluated the overall overlap between all published catalogues, dereplicating the 345,613 sequences from all catalogues into 239,298 species-like vOTUs. Strikingly, 82% of the sequences (n = 195,153), were found to be unique to one catalogue, suggesting low redundancy within these catalogues and justifying the need for a unified resource. The MGV was found to have the largest number of sequences shared with other catalogues, which can be explained by the large size of the resource, and the overlap in the samples included in these mining efforts (**Fig. 1B-C**). Most catalogues contained more than 50% unique sequences that were not found in any other published catalogues, except for the IMG/Vr Gut subset (25% of unique vOTUs) and the KGP (28%). Surprisingly, only 38 vOTUs appeared in six or more of the studies, with one found in all eight adult resources. All 38 vOTUs are bacteriophages belonging to the *Caudoviricetes* Class, potentially infecting genera such as *Bacteroides*, *Bifidobacterium* and *Phocaeicola.* Importantly, 29 of them were found to be integrated in their host genomes, giving a potential explanation for their ubiquitous presence in the catalogues.

### 2. Screening viral sequences from more than 7,000 infant gut metagenomes

To complement the large proportion of healthy adult gut metagenomes previously screened in the published catalogues, we selected 12 large-scale infant studies, with fecal samples collected between birth and 2 years of age from 9 different countries (**Supplemental File 1**). Notably, the largest collection of 2328 samples that derived from the HELMi (Health and Early life Microbiota) cohort dataset (PRJEB70237) also included samples from the infant’s parents (19). A total of 7,867 fecal metagenomes were assembled and screened for viral sequences. We retrieved 1,205,739 putative viral sequences, among which 44,525 high quality and 8,360 complete viral genomes (**Fig. 2A**). It is important to note that our screen used a particularly lenient threshold of viral quality control, in order to allow for a broader retrieval of the viral diversity as additional quality filters were used when merging these putative viral sequences with the other viral resources. These sequences were clustered into 648,848 species-level vOTUs, and as expected, 78% of these vOTUs were singletons, once again highlighting the high viral diversity associated with the human gut.

**Fig 2:**
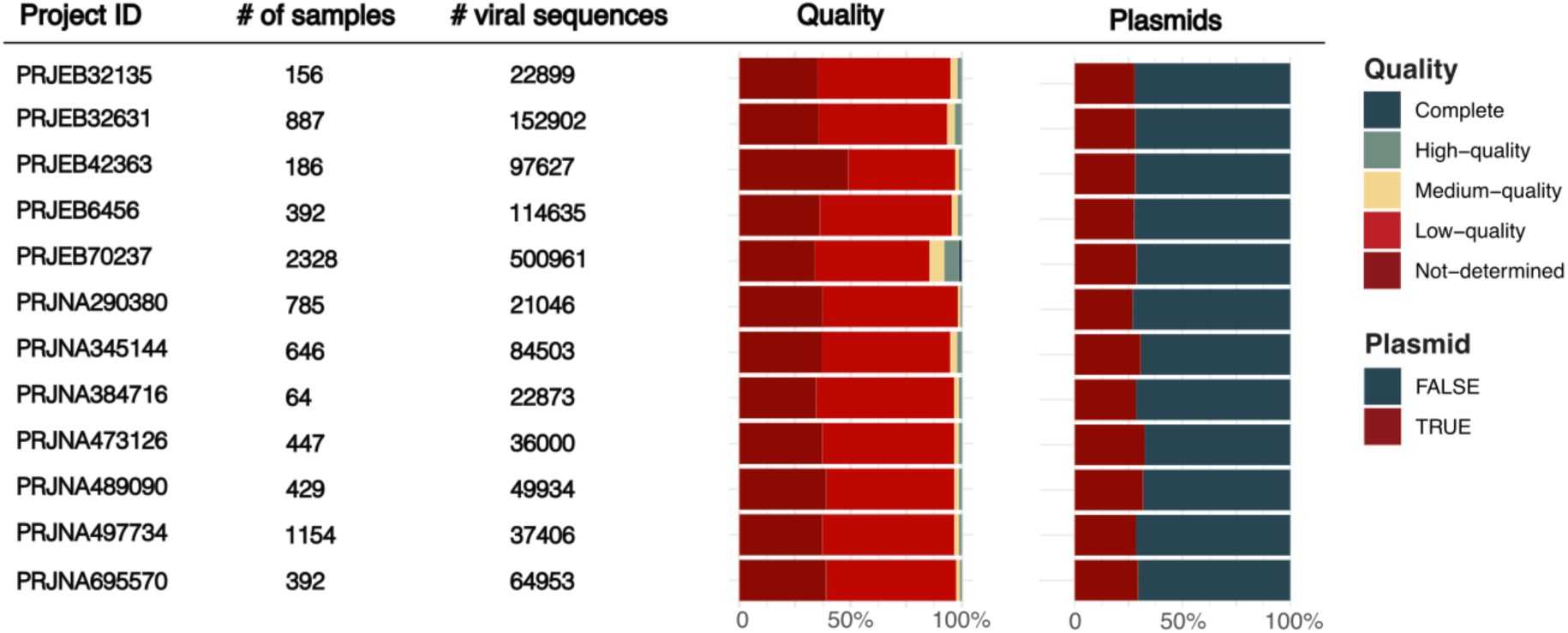
Viral screening of 7,000 infant fecal metagenomes. Overview for each infant project of the number of samples, number of putative viral sequences retrieved and their quality as well as the potential plasmid contamination.

### 3. Building a modular and reusable unified human gut viral resource

From the previously published mining efforts and the additional infant viral sequence collection, we built a unified resource called Aggregated Gut Viral Catalogue (AVrC). After clustering the viral sequences retrieved from the eight published catalogues, the gut subset of the IMG/Vr and the viral sequences retrieved from our infant fecal metagenome screening, we selected vOTUs for which the representative sequence was longer than 5000bp or was annotated as “high-quality” or “Complete” by CheckV. Each putative viral sequence of the catalogue was annotated for sequence quality, potential plasmid contamination, predicted viral taxonomy, predicted viral lifestyle and putative host. The first release of the AVrC contains a total of 1,018,941 unique sequences clustered into 449,859 vOTUs, with 8% complete (n = 36,802), 21% high (n = 93,290) and 22% medium (n = 98,374) quality representative sequences (**Fig. 3A**). Importantly, most vOTU of the AVrC are singleton (65%, n = 294,300), and the vOTU accumulation plot suggests that despite the large-scale efforts in mining the human gut virome, the total species-level viral diversity has not yet been captured (**Fig. 3B).**

**Fig 3:**
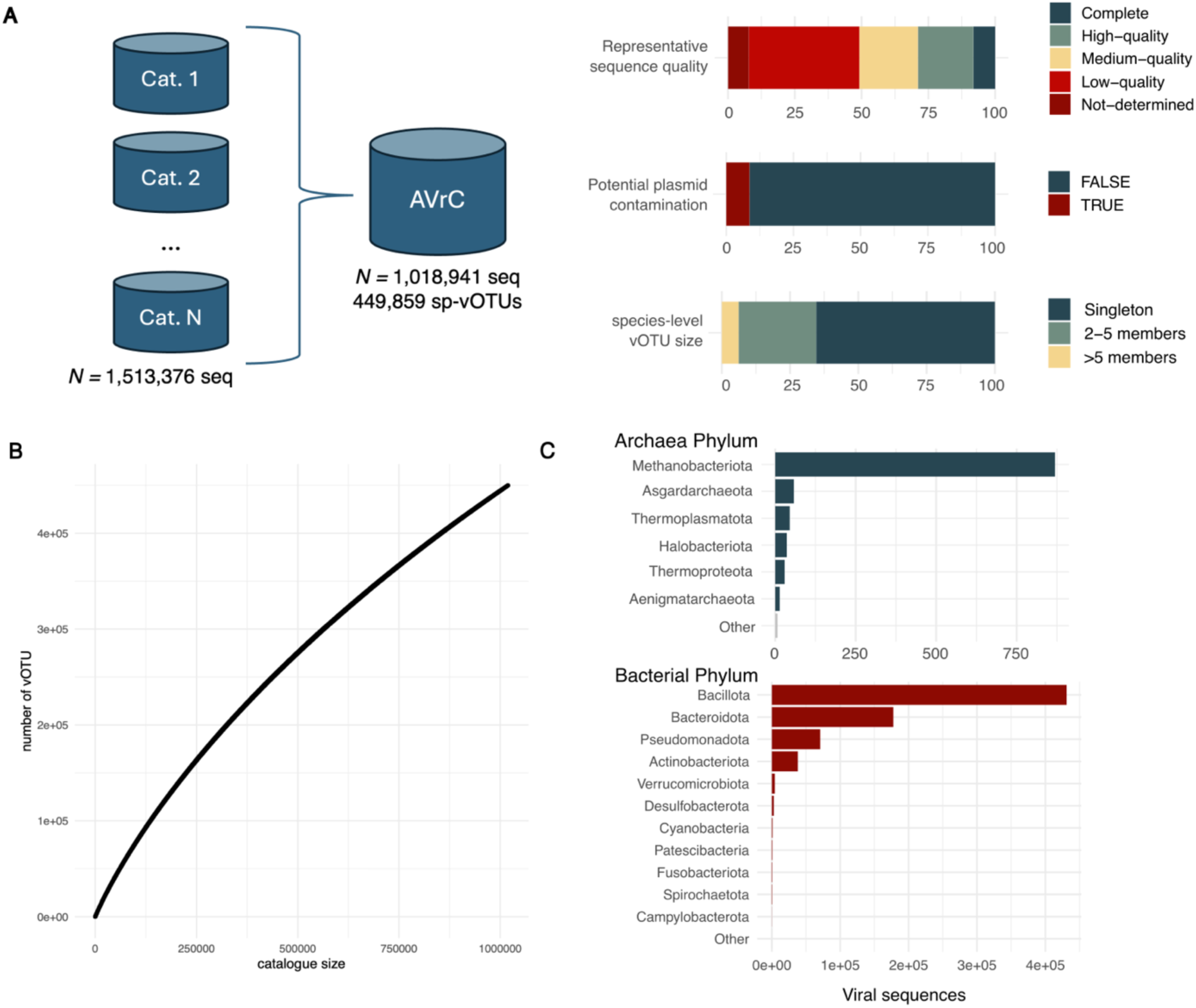
AVrC overview. A: Schematic overview of the AVrC construction. For each vOTU, the representative sequence quality was assessed using CheckV and the potential plasmid contamination was assessed using geNomad. The vOTU size was calculated as the number of sequences grouped into a single cluster by mmSeqs2. B: Accumulation curves of the AVrC at the species-level vOTU. C: Predicted host phylum distribution for the viral sequences contained in the AVrC. The putative host for each viral sequence was obtained from iPHoP. Sequences without any predicted putative host are not displayed in the figure.

As expected, the three most abundant viral classes retrieved are the bacteriophage classes with *Caudoviricetes* constituting 83% (n = 373,751), *Malgrandaviricetes* making 2% (n = 8,083) and *Faserviricetes* being 0.1% (n = 847) of the catalogue’s vOTUs. We estimated that the AVrC contains 58% of temperate bacteriophages (n = 263,079), and 34% of virulent bacteriophages (n = 153,738), with the remainder vOTUs lifestyle being uncertain. Using iPhop, 30,5179 (68%) of the vOTUs could be associated to at least one predicted prokaryotic host, and the predicted hosts for these viral species were part of the Bacillota (45% n = 205,263), Bacteroidota (17%, n = 76,502); Pseudomonadota (8%, n = 33,909) and Actinobacteriota (4%; n = 17,680) phylum, corresponding to the major bacterial taxa found in the human gut (**Fig. 3C**). The number of vOTU retrieved per phylum was strongly correlated to the number of predicted host genera per phylum (Spearman correlation, p< 2.2e-16, rho= 0.96), suggesting that the taxonomic diversity of these phylum in the gut could be driving the high representation of vOTU infecting these taxa in the AVrC. Interestingly, the catalogue also included phages infecting less common prokaryotic groups; as an example, the AVrC contains 393 vOTUs predicted to infect Spirochaetota species found exclusively in non-industrialized population (20) and contains 610 vOTUs predicted to infect archaeal species.

Importantly, the AVrC was implemented as a modular relational database to ensure easy additions or updates of the datasets and annotations. Indeed, as new catalogues and new bioinformatic tools are updated and published, we anticipate a need to continuously update the resource. The database is composed of sequence files in Fasta format containing all the sequences or the subset of representative sequences of each vOTU. All sequences can be easily linked to their individual annotations by five different tools (CheckV (21), geNomad (22), PhaGCN (23), PhaTyp (24) and iPHoP (25)). Finally, these tool annotations were combined to generate 3 global annotation tables that summarize the sequence quality, viral taxonomy and lifestyle, and host information (**Fig. 4A**).

**Fig 4:**
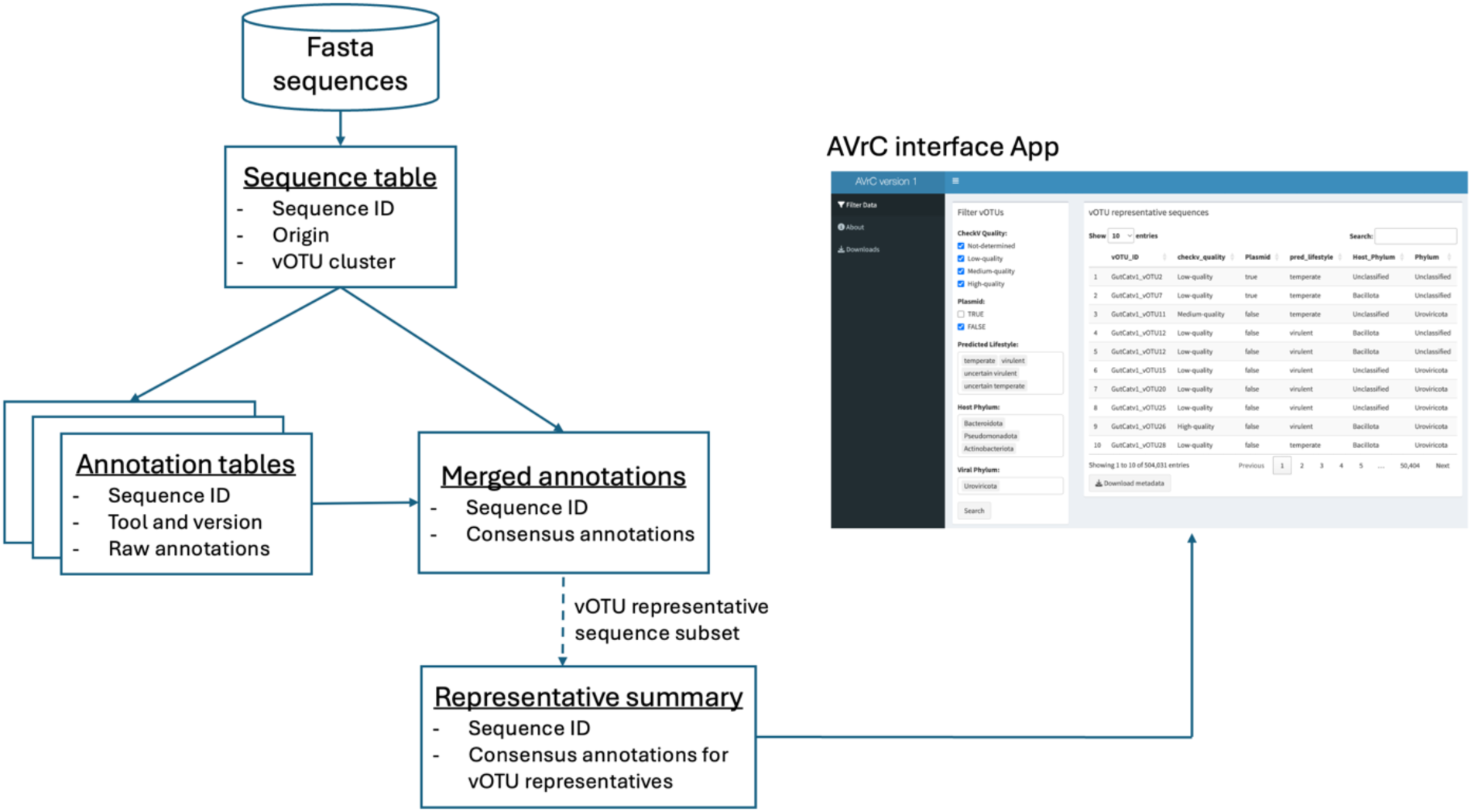
AVrC structure and interface. AVrC database schematical structure. The AVrC database contains a fasta sequence catalogue containing the viral sequences in a Fasta format. The annotations of the sequences are grouped in three types of tables: (1) the raw output of each annotation tools, (2) the merged and harmonized annotations recapitulating the information concerning the sequence’s quality, taxonomy and lifestyles and the predicted host information, and (3) a summary table containing the merged information for the vOTU representative sequences. This summary table is searchable through the AVrC interface, allowing users to select and search and select subsets of the dataset without any prior coding experience.

To enable a greater reusability of the AVrC, the viral sequence catalogue and the viral annotations are available as a Zenodo archive and through the AVrC website (https://github.com/aponsero/Aggregated_Viral_Catalogue). The AVrC website enables users to quickly select and download subsets of the AVrC catalogue through a user-friendly interface. The AVrC search interface allows users to select vOTU representative sequences according to their sequence quality, length, viral taxonomy and putative hosts, and download the sequences a summary of the annotations (**Fig. 4B**). Data subsets of interest are also provided for an easy direct download of the sequences and annotations, including a subset containing only high-quality vOTU representative sequences and the subset of all bacteriophage sequences.

## Discussion

Recently, several large-scale efforts have been made to better understand and characterize the human gut virome, in particular by mining human gut metagenomes to generate viral catalogues. These resources are critical to enable the description of the diversity of viruses in human associated ecosystems and constitute valuable resources for the research community. Typically, each of these studies leveraged different tools, methods, and applied different quality control criteria, making the direct comparison of these resources impossible.

Here, we surveyed eight previously published studies and the subset of human gut viral sequences from the IMG/Vr database (10–18). As expected, a large proportion of the fecal metagenomes surveyed by these catalogues were collected from healthy adults from western countries, reflecting the current known bias in the human metagenome sampling. When assessing the sequence quality of these previously published studies, we observed a high prevalence of potential plasmid sequences in the datasets, in particular for the oldest resources. We therefore suggest that future viral mining efforts include a quality control step for potential plasmid contamination (Camargo, Roux, *et al.* 2023). Strikingly, most of the viral sequences were found to be unique to one catalogue, even though 30% of the metagenomes were mined by at least 2 studies. This result could be explained by the impact of the computational viral detection methods on the viral sequences retrieved from these metagenomes, as suggested by the benchmark of these computational phage detection approaches (26).

To complement the previous mining efforts, we screened an additional 7,867 fecal infant gut metagenomes for viral sequences, retrieving over 1.2 million putative viral sequences that were clustered into 648,848 species-level viral operational taxonomic units (vOTUs). These infant viral sequences were combined with previously published human gut viral catalogues to construct the Aggregated Gut Viral Catalogue (AVrC) - a unified modular resource containing 1,018,941 dereplicated viral sequences clustered into 449,859 vOTUs. Despite these large-scale mining efforts, our clustering results suggest that the species-level viral diversity in the human gut has not been completely captured.

The AVrC was constructed as a modular relational database that provides extensive annotations of the sequences included in the catalogue, including the sequence quality and potential plasmid contamination, viral taxonomy, predicted lifestyle, and putative microbial hosts. The AVrC is available through a user-friendly web interface to enable easier customized querying and retrieval of sequences/annotations. In the future, we aim to expand the AVrC through the continued integration of novel catalogues recently published. As an example, a novel catalogue surveying infant fecal viral diversity (27), and a virome catalogue of a large colorectal cancer screening program ((28)), were published during the redaction of this manuscript and will be integrated in the upcoming version 2 release of the AVrC. Additionally, recent reports showed the potential of long-read sequencing technology and hybrid approaches in the exploration of the human gut virome (29). Finally, as computational methods to annotate and explore the viral diversity are in constant improvement, the AVrC was built to rapidly allow for the addition of novel annotations and to facilitate the update of the annotations when new reference databases are published.

Importantly, at the time of writing, other viral catalogue unification efforts are underway, in particular the Unified Human gut Virome Catalog (UHGV, available at https://github.com/snayfach/UHGV). However, the structure and aims of the AVrC and UHGC are distinct, and we believe the two resources to be complementary. Indeed, the UHGC aims to unify previously published gut viral catalogue and annotate the high-quality viral sequences using a novel and curated computational pipeline to generate a high-quality viral catalogue. On the other hand, the AVrC is designed to be an aggregation of resources, more modular and that leaves the user free to choose subsets of the catalogue depending on their needs. Both approaches are addressing a current need to harmonize large scale viral catalogues into a consistent dataset.

## Methods

### Dada source: Published gut viral catalogues and infant gut metagenomes viral mining

Literature search for previously published human gut viral mining efforts from PubMed was performed in 2023 and allowed the identification of eight relevant studies (**Table 1**). The IMG/Vr and CHVD catalogues were filtered to retrieve only viral sequences obtained from human fecal metagenomes. The original sequence names were mapped to a unified naming convention across the datasets for easier integration in the AVrC. The mapping between the original and new sequence names is available in the Zenodo archive and on the AVrC website.

We additionally selected 12 gut metagenome projects from large-scale infant birth cohorts and downloaded the metagenomes directly from NCBI or ENA. Quality controlled reads were assembled using Megahit v1.2.9 (30) for PRJEB70237 and the METAnnotatorX2 pipeline (31) using Spades v3.15 (32) (other projects). Assembled contigs longer than 500bp and with a coverage above 5x were classified as viral or non-viral using DeepVirFinder v1.0 (4) and VirSorter2 v2.2.3 (6). Putative viral sequences were defined as follows: DeepVirFinder score above 0.9 or VirSorter2 viral/prophage classification. The putative viral contigs were further confirmed using CheckV v0.8.1 (21) and contigs longer than 1kb with no detected viral genes and at least one cellular gene was discarded. The sequences were dereplicated using MMseqs2 (33) with 99% identity over 90% of shortest sequences.

### Viral sequence clustering and annotations

The putative viral sequences were clustered into species-like vOTU using MMseqs2 (33) with a 95% identity over 75% of shortest sequences as commonly used (34). The longest sequence for each cluster was chosen as a vOTU representative. Finally, the vOTUs with a representative sequence above 5,000 bp or classified as “high-quality” or “Complete” by CheckV were selected and kept in the AVrC.

All sequences were annotated by geNomad v1.7.4 (22) and PhaGCN (23) to obtain high level viral classification, following the new ICTV convention. GeNomad was also used to identify potential plasmid sequences. The putative viral lifestyle strategy was determined using PhaTyp (24) as well as the annotations derived from CheckV and geNomad. Briefly, four categories were generated: temperate (PhaTyp classifies the sequence as temperate with a score of >= 0.7 or CheckV and geNomad predict a prophage sequence), uncertain temperate (PhaTyp classifies the sequence as temperate with a score of < 0.7), virulent (PhaTyp classifies the sequence as virulent with a score of >= 0.7) and uncertain virulent (PhaTyp classifies the sequence as virulent with a score of < 0.7). Host prediction for the viral sequences was obtained using iPHOP v1.3.3 (25), a tool that leverages six distinct methods including both host-based tools (e.g. CRISPR markers, prophage in host genome, etc.) and phage-based tools (e.g. alignment with phages with known hosts) and merges their results to provide the user with a candidate host genus for each viral sequence.

### AVrC database and interface

The AVrC sequence catalogue and annotations are available in fasta and csv format in a Zenodo archive (10.5281/zenodo.11426065). The AVrC website interface was created using the ShinyR v1.8.1 package, and the ggplot2 v3.5.0 for graphical representations. All major browsers support access to our database website, accessible at https://github.com/aponsero/Aggregated_Viral_Catalogue.

## Data availability statement

The data that support the findings of this study are in Zenodo DOI 10.5281/zenodo.11426065, the github repository (https://github.com/aponsero/Aggregated_Viral_Catalogue) and in the Supplementary material.

## Acknowledgments

We thank the Finnish IT Centre for Science and the NBI Research computing for providing the computational resources used for this project. We thank the other group members of the Microbes inside lab for their helpful contributions, in particular Roosa Jokela and Dollwin Matharu. We thank Prof. Kaija-Leena Kolho for helpful discussions. Finally, we thank Dr. Andrea Telatin for his support in finalizing this study and the QIB core bioinformatics team.

## Supporting information captions

**Suplementary File 1: Description of the samples screened by the gut viral catalogues.**

